# COP1 destabilizes DELLA proteins in *Arabidopsis*

**DOI:** 10.1101/2020.01.09.897157

**Authors:** Noel Blanco-Touriñán, Martina Legris, Eugenio G. Minguet, Cecilia Costigliolo-Rojas, María A. Nohales, Elisa Iniesto, Marta García-León, Manuel Pacín, Nicole Heucken, Tim Blomeier, Antonella Locascio, Martin Černý, David Esteve-Bruna, Mónica Díez-Díaz, Břetislav Brzobohatý, Henning Frerigmann, Matías D. Zurbriggen, Steve A. Kay, Vicente Rubio, Miguel A. Blázquez, Jorge J. Casal, David Alabadí

## Abstract

DELLA transcriptional regulators are central components in the control of plant body form in response to the environment. This is considered to be mediated by changes in the metabolism of the hormones gibberellins (GAs), which promote the degradation of DELLAs. However, here we show that warm temperature or shade reduced the stability of a GA-insensitive DELLA allele in *Arabidopsis*. Furthermore, the degradation of DELLA induced by the warmth anticipated changes in GA levels and depended on the E3 ubiquitin ligase CONSTITUTIVELY PHOTOMORPHOGENIC1 (COP1). COP1 enhanced the degradation of normal and GA-insensitive DELLA alleles when co-expressed in *N. benthamiana.* DELLA proteins physically interacted with COP1 in yeast, mammalian and plant cells. This interaction was enhanced by the COP1 complex partner SUPRESSOR OF *phyA-105* 1 (SPA1). The level of ubiquitination of DELLA was enhanced by COP1 and COP1 ubiquitinated DELLA proteins in vitro. We propose that DELLAs are destabilized not only by the canonical GA-dependent pathway but also by COP1 and that this control is relevant for growth responses to shade and warm temperature.

**Significance:** DELLA proteins are plant-specific transcriptional regulators that act as signaling hubs at the interface between the environment and the transcriptional networks that control growth. DELLAs are destabilized by the growth-promoting hormone gibberellin, whose levels are very sensitive to environmental changes. Here we describe an alternative pathway to destabilize these proteins. We show that DELLAs are substrate of COP1, an E3 ubiquitin ligase that increases its nuclear activity to promote growth in response to shade or warmth. Our results also show that the destabilization of DELLAs by COP1 precedes the action of gibberellins, suggesting the existence of a sequential mechanism to control the stability of these proteins.

---

A plant can adopt markedly different morphologies depending on the environment it has to cope with. This plastic behavior relies on highly interconnected signaling pathways, which offer multiple points of control (1). Light and temperature are among the most influential variables of the environment in plant life. For instance, light cues from neighboring vegetation as well as elevated ambient temperature (e.g. 29 °C) enhance the growth of the hypocotyl (among other responses) respectively to avoid shade (2) and enhance cooling (3).

Several features place DELLA proteins as central elements in environmental responses (4). First, DELLAs are nuclear-localized proteins that interact with multiple transcription factors and modulate their activity (5). Second, they are negative elements in the gibberellin (GA) signaling pathway and their stability is severely diminished upon recognition of their N-terminal domain by the GA-activated GIBBERELLIN INSENSITIVE1 (GID1) receptor, which recruits the SCF^SLY1/GID2^ complex to promote their ubiquitination-dependent degradation by the proteasome (6). Third, GA metabolism is tightly regulated by the environment, for instance, shade and warm temperature induce GA accumulation (3, 7).

DELLA levels increase during illumination of dark-grown seedlings and promote transcriptional changes associated with photomorphogenesis (8, 9). On the contrary, they decrease during the night and in response to shade inflicted by neighbor plants or to warm ambient temperature, allowing the promotion of hypocotyl and/or petiole elongation by transcription factors such as PHYTOCHROME INTERACTING FACTOR4 (10–13). Interestingly, the role of DELLAs in all these processes is the opposite to that of CONSTITUTIVELY PHOTOMORPHOGENIC1 (COP1), another central regulator of light and temperature responses. COP1 is an E3 ubiquitin ligase that promotes proteasome-dependent degradation of a number of transcription factors involved in light and temperature signaling, and becomes inactivated by light perceived by phytochromes and cryptochromes and by low-to-moderate temperature (14–19). It requires the activity of the SUPRESSOR OF *phyA-105* proteins (SPA1 to 4 in *Arabidopsis*) to be active in vivo (20). Here we show the direct physical interaction between DELLAs and COP1/SPA1 complex and propose a mechanism of regulation of DELLA stability different from the canonical GA signaling pathway.

## Results

### Warm temperature or shade decrease the abundance of a GA-resistant DELLA protein

Warm temperatures or shade decrease the abundance of the DELLA protein REPRESSOR OF *ga1-3* (RGA) (Fig. 1*A*) (10, 13). Two observations indicate that changes in GA cannot fully account for these responses. First, increasing doses of the GA-inhibitor paclobutrazol (PAC) elevated RGA nuclear abundance observed by confocal microscopy in a *pRGA:GFP-RGA* line (21), but the responses to shade or warmth persisted even under saturating levels of the inhibitor (Fig. 1*A*). Second, warm temperature or shade reduced the levels of rga-Δ17, a mutant version of RGA that is fully resistant to GA, in the *pRGA:GFP-(rga-Δ17)* line (Fig. 1 B and C) (22). Changes in *RGA* transcript levels do not mediate the RGA abundance response to shade (13) or warm temperature (*SI Appendix*, Fig. S1). Importantly, RGA abundance responses were fully impaired by treatment with the inhibitor of the 26S proteasome MG132 (Fig. 2*A*). Altogether, these results suggest the existence of non-canonical pathway of DELLA degradation.

**Fig. 1.**
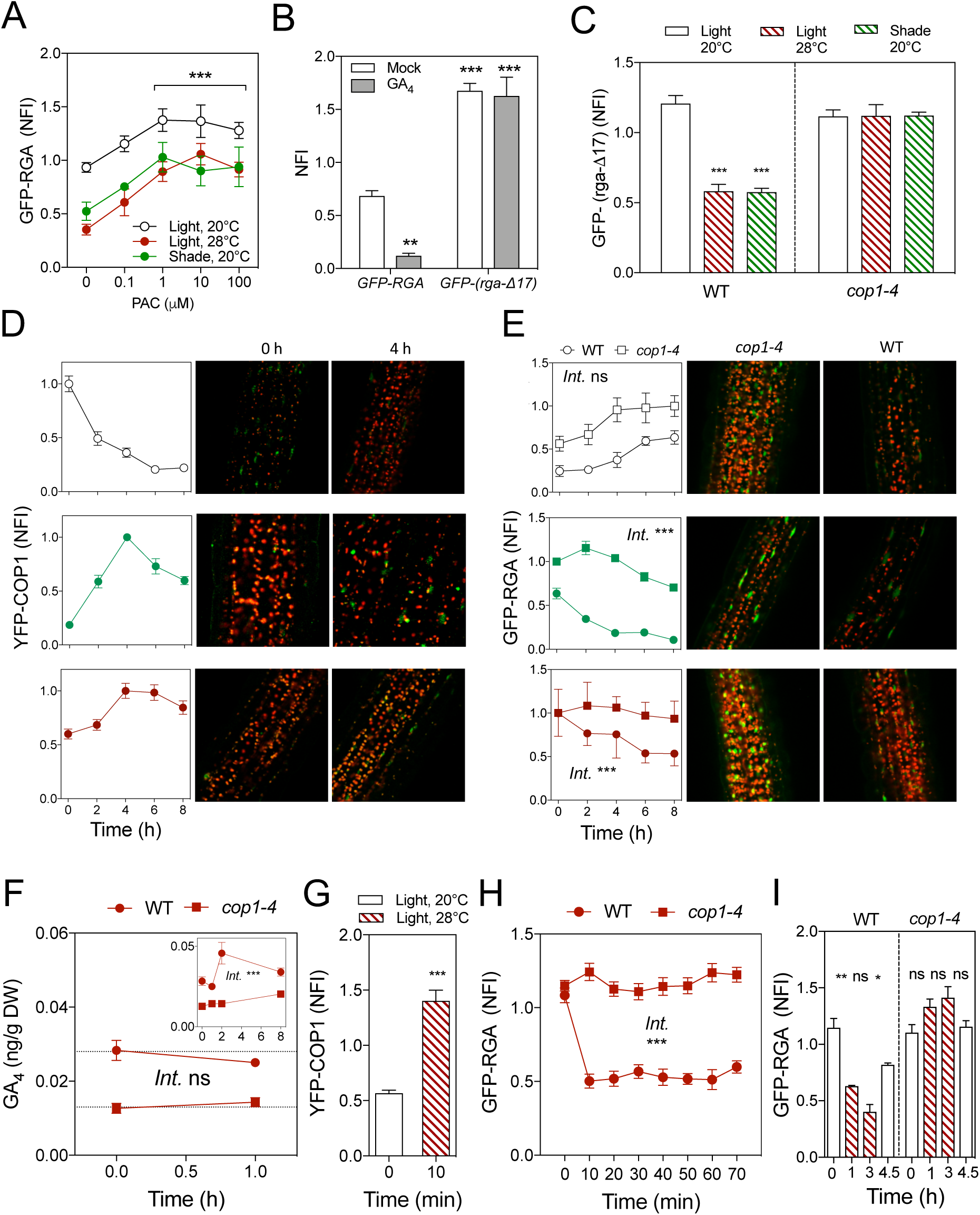
COP1 regulates RGA levels in *Arabidopsis* hypocotyls in response to changes in light and temperature. (A) GFP-RGA levels respond to changes in light and temperature independently of the GA pathway. Seedlings were treated with PAC for 8 h before subjected to shade or 29 °C for 4 h. The plot shows the normalized fluorescence intensity (NFI) in nuclei. Data are average of 5-10 seedlings (10-40 nuclei were averaged per seedling replicate). (B) GFP-(rga-Δ17) is resistant to GA-induced degradation. Seedlings of *pRGA:GFP-RGA* and *pRGA:GFP-(rga-Δ17)* lines were mock-treated or treated with 3 μM GA_4_ for 4 h. The plot shows the NFI in nuclei. Data are average of 6-9 seedlings (10-27 nuclei were averaged per seedling replicate). (C) GFP-(rga-Δ17) levels are reduced in response to shade or 29 °C in a COP1-dependent-manner. NFI data are means and SE of 6-14 seedlings (10-30 nuclei were averaged per seedling replicate). (D-E) Dynamics of nuclear accumulation of YFP-COP1 (D) in the wild-type and of GFP-RGA (E) in the wild-type (circles) and *cop1-4* mutants (squares) after the transfer from darkness to light (white symbols), light to shade (green symbols), and 22 °C to 29 °C (red symbols). NFI data are means and SE of 18 seedlings (30-50 nuclei were averaged per seedling replicate). In (E), the slopes are significantly different (*P* <0.001) between wild-type and *cop1* seedlings transferred to shade (first 4 h) or to 29 °C but not in seedlings transferred from darkness to light. Images of representative hypocotyls from wild-type and *cop1-4* were taken by confocal microscopy at the 8 h time point. (F) Time course of GA_4_ levels in wild-type and *cop1-4* seedlings before (time =0 h) and after transfer to 29 °C. Data are means and SE of three independent biological replicates and the inset shows an extended time-course. (G) Nuclear levels of YFP-COP1 in seedlings maintained at 22 °C or after transfer to 28 °C for 10 min. NFI data are average of 11-12 seedlings (10-20 nuclei were averaged per seedling replicate). (H) Time course of nuclear levels of GFP-RGA in wild-type and *cop1-4* seedlings before (time =0 h) and after transfer to 29 °C. NFI data are means and SE of 6-13 seedlings (10-35 nuclei were averaged per seedling replicate). (I) The reduction of GFP-RGA levels in response to 29 °C is reversible and dependent on COP1. Seedlings of *pRGA:GFP-RGA* in the WT and *cop1-4* backgrounds were returned to 22 °C after 3 h-treatment of 29 °C. NFI data are means and SE of 18 seedlings (10-40 nuclei were averaged per seedling replicate). Asterisks in A, B, C, G and I indicate that the difference is statistically significant (Student’s *t* test, * *P* < 0.05, ** *P* < 0.01, *** *P* < 0.005; ns, non-significant). In I, asterisks and ns refer to the comparison between adjoining bars. Asterisks in E, F and H indicate that the term accounting for the interaction between temperature and the *cop1* mutation is statistically significant at *P* < 0.0005 in multiple regression analysis (ns, non-significant).

**Fig. 2.**
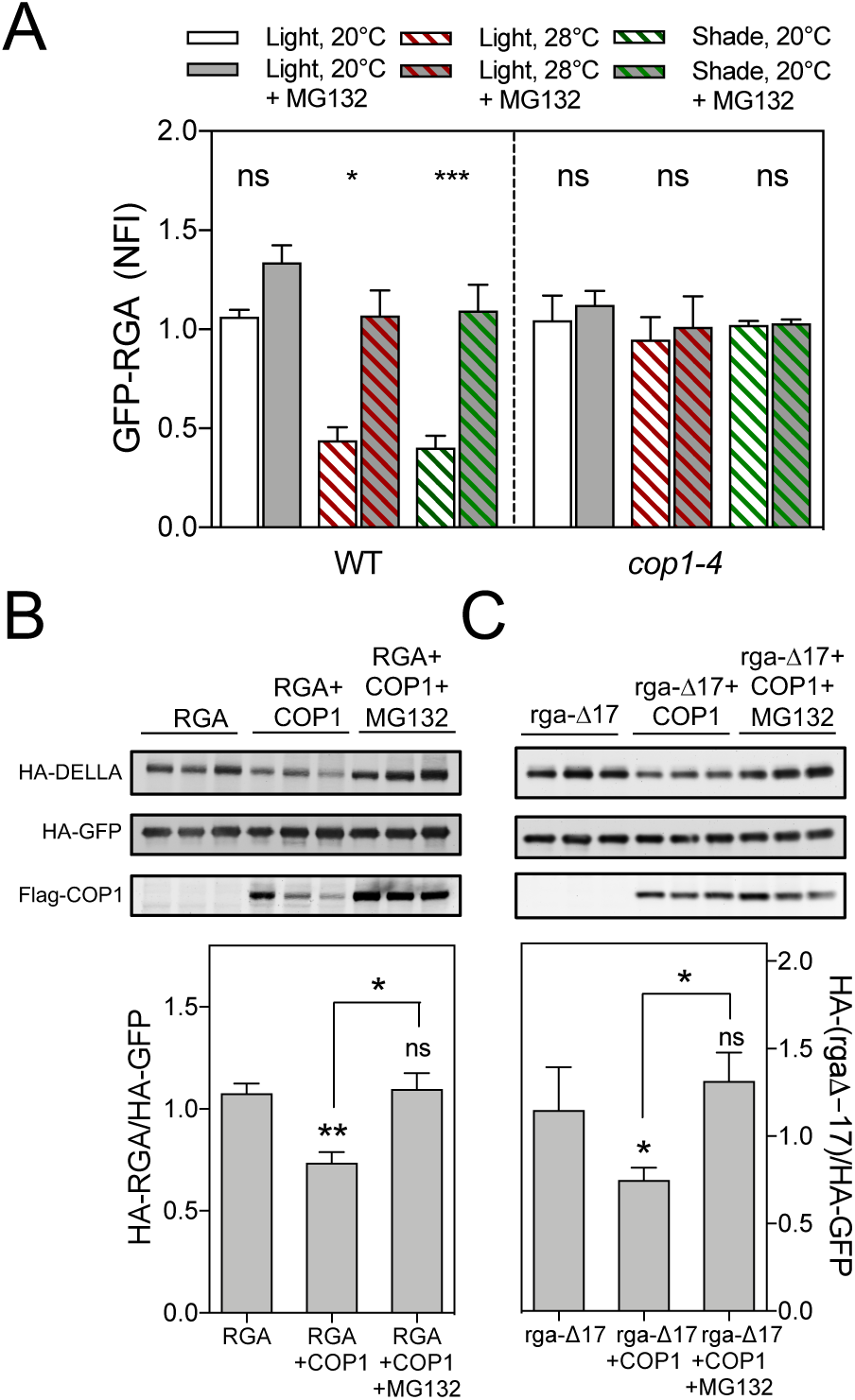
COP1 destabilizes DELLAs. (A) The reduction of GFP-RGA levels by warm temperature or shade requires the 26S proteasome and COP1. NFI data are means and SE of 6-9 seedlings (10-30 nuclei were averaged per seedling replicate). Asterisks indicate that the difference is statistically significant (Student’s *t* test, * *P* < 0.05 and *** *P* < 0.001; ns, non-significant). (B-C) COP1 destabilizes RGA (B) and the GA-resistant rga-Δ17 (C) in *N. benthamiana* leaves. HA-RGA and HA-(rga-Δ17) were transiently expressed alone or with FLAG-COP1 in leaves of *N. benthamiana*. For MG132 treatments leaves were infiltrated with a solution of 25 μM of the inhibitor 8 h before sampling. HA-GFP was used as control to demonstrate the specificity of COP1 action. Blots show data from three individual infiltrated leaves per mixture. Plots show HA-RGA and HA-(rga-Δ17) normalized against HA-GFP. Data are means and SE of 3 leaves from one experiment, repeated twice with similar results. Asterisks indicate that the difference is statistically significant (Student’s *t* test, * *P* < 0.05 and ** *P* < 0.01; ns, non-significant).

### COP1 affects RGA before modifying GA levels

RGA levels are elevated in *cop1-4* seedlings (23). Compared to light at moderate temperature, darkness, shade or warm temperature increased the nuclear abundance of COP1 (14, 24, 25) in a *35S:YFP-COP1 cop1-4* line (26), while reducing RGA levels (9, 10, 13) (Fig. 1 D and E). The light-induced increase in RGA showed wild-type kinetics in the *cop1-4* seedlings (note parallel curves), suggesting that this response is driven by a COP1-independent light-induced down-regulation of GA biosynthesis (8, 9, 27, 28). Conversely, *cop1-4* seedlings grown in the light at moderate temperature (20 °C) and transferred either to shade at the same temperature or to light at 29 °C, showed a markedly delayed decrease in GFP-RGA (Fig. 1*E*). GA_4_ levels were unaffected by transferring the seedlings from 20 °C to 29 °C for 1 h (Fig. 1*F*), whilst 10 min of warm temperature were enough to drive significant nuclear accumulation of COP1 (Fig. 1*G*). Warm temperature decreased GFP-RGA levels with a lag of less than 10 min; this response was absent in the *cop1-4* mutant and reversible after the return to 20 °C (Fig. 1 H and I). Warm temperature did increase GA levels, but only after 2 h (Fig. 1*F*, inset). In the *cop1-4* mutant the level of GA was reduced, as reported for pea (29), and failed to respond to temperature (Fig. 1*F*). Taken together, these results indicate that the normal shade-or warmth-induced degradation of RGA requires COP1 and anticipates changes in GA.

### COP1 promotes degradation of a GA-resistant DELLA protein

In *Arabidopsis*, warm temperature or shade failed to reduce the nuclear abundance of RGA in the presence of the 26S proteasome inhibitor MG132 and/or the *cop1-4* mutation (Fig. 2*A*). The degradation of the rga-Δ17 also required COP1 (Fig. 1*C*). Co-expression of COP1 caused 26S proteasome-dependent decrease of HA-RGA and also of HA-(rga-Δ17) in leaves of long-day grown *Nicotiana benthamiana* plants, while it had no impact on levels of the unrelated protein HA-GFP (Fig. 2 B and C). Warm temperature decreased HA-(rga-Δ17) in a COP1-mediated manner (*SI Appendix*, Fig. S2). This suggests that COP1 mediates the destabilization of RGA by non-canonical mechanisms. To explore this possibility, we first investigated whether COP1 physically interacts with DELLA proteins.

### COP1 interacts physically with GAI and RGA in yeast

We performed yeast two-hybrid (Y2H) assays between COP1 and the two DELLAs with a major role in light- and temperature-dependent growth, RGA and GIBBERELLIC ACID INSENSITIVE (GAI) (10, 11, 30). To avoid the reported strong auto-activation of full-length DELLAs in yeast, we used N-terminal deleted versions named M5GAI and RGA52 (12, 31). COP1 was able to interact with both (Fig. 3*A*). SUPRESSOR OF *phyA-105* 1 (SPA1) and other SPA proteins involved in a functional complex with COP1 (20, 32) were also able to interact with GAI and RGA in Y2H assays (Fig. 3*B*).

**Fig. 3.**
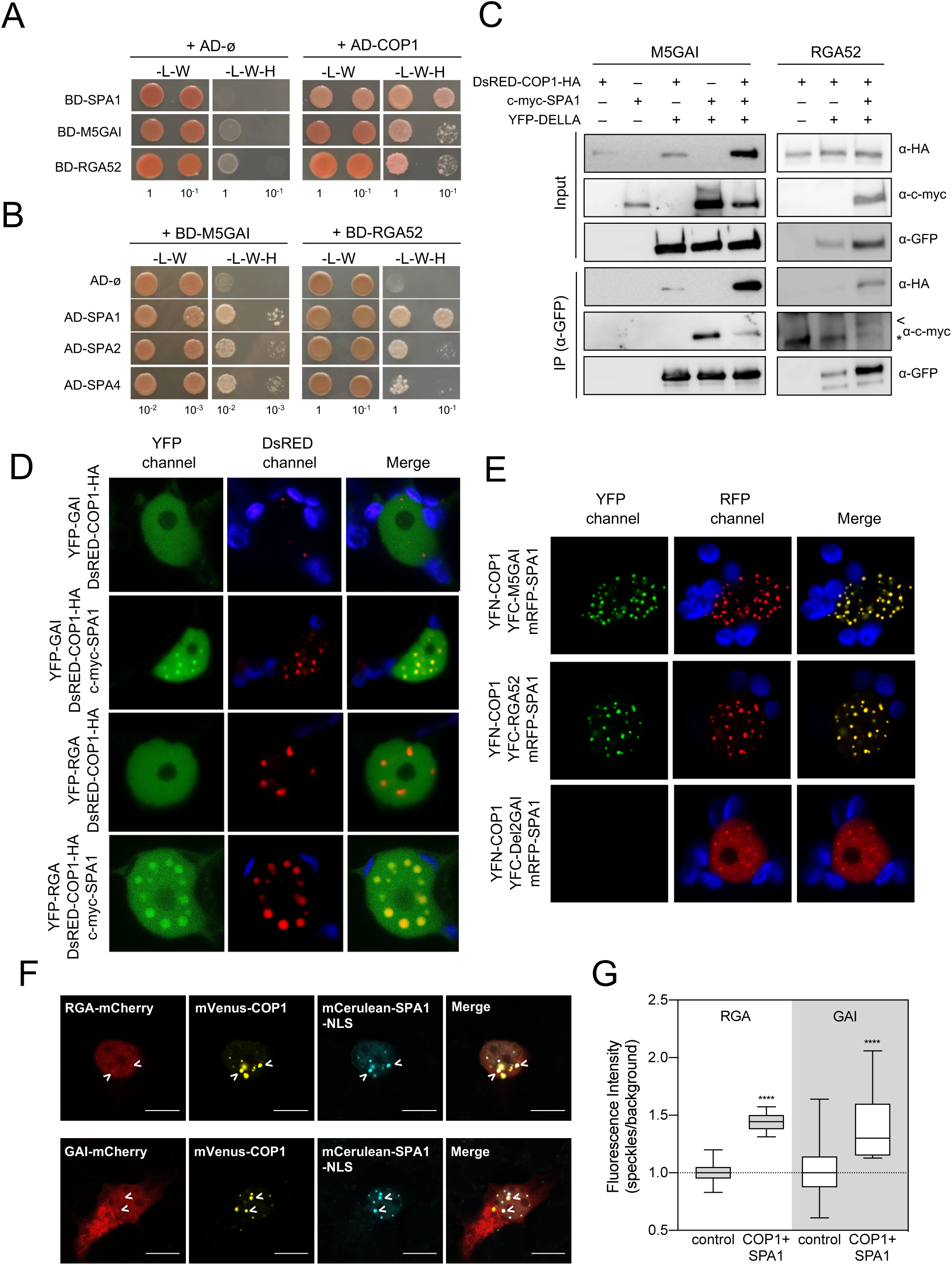
COP1 physically interacts with DELLA proteins. (A-B) Y2H assays showing the interaction between N-terminal deleted versions of GAI and RGA with COP1 (A) and SPAs (B). L, leucine; W, tryptophan; H, histidine. Numbers indicate the dilutions used in the drop assay. (C) Co-immunoprecipitation assays showing interactions in planta. YFP-M5GAI and YFP-RGA52 were transiently expressed in leaves of *N. benthamiana* together with DsRED-COP1-HA, c-myc-SPA1 or both. Proteins were immunoprecipitated with anti-GFP antibody-coated paramagnetic beads. Leaves expressing DsRED-COP1-HA or c-myc-SPA1 alone were used as negative controls. The arrowhead and asterisk mark the co-immunoprecipitated c-myc-SPA1 and a non-specific band, respectively. (D) YFP-GAI and YFP-RGA co-localize with DsRED-COP1-HA in nuclear bodies in the presence of c-myc-SPA1. Fusion proteins were transiently expressed in leaves of *N. benthamiana* and observed by confocal microscopy. One representative nucleus is shown. (E) BiFC assay showing that COP1 and SPA1 form a complex with M5GAI or RGA52 in nuclear bodies. The indicated proteins were expressed in leaves of *N. benthamiana* and observed by confocal microscopy. One representative nucleus is shown. (F) Representative HEK-293T cells co-transfected with with mVenus-COP1, NLS-mCerulean-SPA1 and either GAI-mCherry or RGA-mCherry. Scale bar represents 10 μm. The arrowheads point to two representative speckles co-occupied by DELLAs, SPA1 and COP1. (G) Fluorescence intensities of GAI-mCherry and RGA-mCherry in control cells and in cells co-expressing mVenus-COP1 and mCerulean-SPA1-NLS from 10-13 transfected cells. Asterisks indicate that the difference is statistically significant (Student’s *t* test, **** *P* < 0.0001).

### COP1 interacts with GAI and RGA *in planta*

To investigate whether the interaction between DELLAs and COP1 also occurs in plant cells, we first performed co-immunoprecipitation assays in leaves of *N. benthamiana* co-expressing DsRED-COP1-HA and YFP-M5GAI or YFP-RGA52. While DsRED-COP1-HA was pulled down by anti-GFP antibodies from leaf extracts co-expressing YFP-M5GAI and the interaction appeared to be enhanced in the presence of c-myc-SPA1, the DsRED-COP1-HA and YFP-RGA52 interaction was only observed when the three proteins were co-expressed in same leaves (Fig. 3*C*). c-myc-SPA1 was also specifically co-immunoprecipitated with YFP-M5GAI (Fig. 3*C*). These results suggest that SPA1 enhances the interaction between COP1 and DELLA proteins. Consistent with this idea, we observed re-localization of YFP-GAI, YFP-RGA, and RGA52-YFP to nuclear bodies co-occupied by DsRED-COP1-HA in the presence of c-myc-SPA1 (Fig. 3*D* and *SI Appendix*, Fig. S3*A*).

### COP1-SPA1 forms a ternary complex with DELLA

The formation of a ternary complex was evidenced by bi-molecular fluorescence complementation (BiFC) assays in leaves of *N. benthamiana*, in which the co-localization of signals from mRFP-SPA1 and the reconstituted YFP, due to the interaction between YFC-DELLAs and YFN-COP1, was evident in nuclear bodies (Fig. 3*E* and *SI Appendix*, Fig. S3*B*). Similarly, YFP signal in nuclear bodies was observed by co-expressing c-myc-SPA1 (*SI Appendix*, Fig. S3*C*). However, no YFP fluorescence was detected in the absence of SPA1 or when YFC was fused to Del2GAI, a truncated version of GAI that does not interact with SPA1 (Fig. 3*E* and *SI Appendix*, Fig. S3 *B*-*E*). As expected, mRFP-SPA1 was recruited to nuclear bodies when co-expressed with YFN-COP1 (Fig. 3*E* and *SI Appendix*, Fig. S3*B*).

To quantify the interaction between GAI or RGA and the COP1-SPA1 complex we expressed these proteins tagged to fluorescent reporters in mammalian cells. This orthogonal system allows to perform such studies with the components of interest, in the absence of other plant proteins that might interfere with the evaluation. The fluorescence driven by DELLA proteins in the cytosol and nucleus was relatively homogeneous when either GAI or RGA were expressed alone (note the ratio of fluorescence between different nuclear ROIs close to 1, Fig. 3*G*). However, the ratio between DELLA fluorescence inside / outside the speckle-like structures formed in the nucleus by the COP1-SPA1 complex was above 1 (Fig. 3 F and G and *SI Appendix*, Fig. S4), indicating that the COP1-SPA1 complex drags RGA and GAI to the speckles by physical interaction. Taken together, these observations demonstrate that the COP1-SPA1 complex interacts with DELLA proteins.

### COP1 ubiquitinates GAI and RGA in vitro

In vivo levels of ubiquitinated GFP-RGA were enhanced by over-expression of COP1 (*SI Appendix*, Fig. S4). To test whether this is the result of the direct interaction between COP1 and DELLAs, we performed an in vitro ubiquitination assay using recombinant MBP-COP1 and 6xHis-M5GAI or 6xHis-RGA52. A slow-migrating band corresponding to the size of Ub-6xHis-M5GAI or Ub-6xHis-RGA52 was observed only when MBP-COP1 and the E2 enzyme were included in the assays (Fig. 4 A and B). The delayed band did not appear, however, when Zn^2+^ ions, which are required for the proper arrangement of the RING domain of E3 ubiquitin ligases like COP1 (33), were excluded from the reaction mixtures (Fig. 4 A and B). To confirm that the slow migration of 6xHis-M5GAI and 6xHis-RGA52 is due to ubiquitination, we repeated the assay for 6xHis-M5GAI in the presence of HA-tagged ubiquitin. We detected low-migrating bands in the immunoblot with anti-GAI antibody when free ubiquitin was included in the assay, which were further upshifted when we used the HA-tagged version of ubiquitin instead (Fig. 4*C*). This result indicates that M5GAI and RGA52 are targets of the E3 ubiquitin ligase activity of COP1 in vitro.

**Fig. 4.**
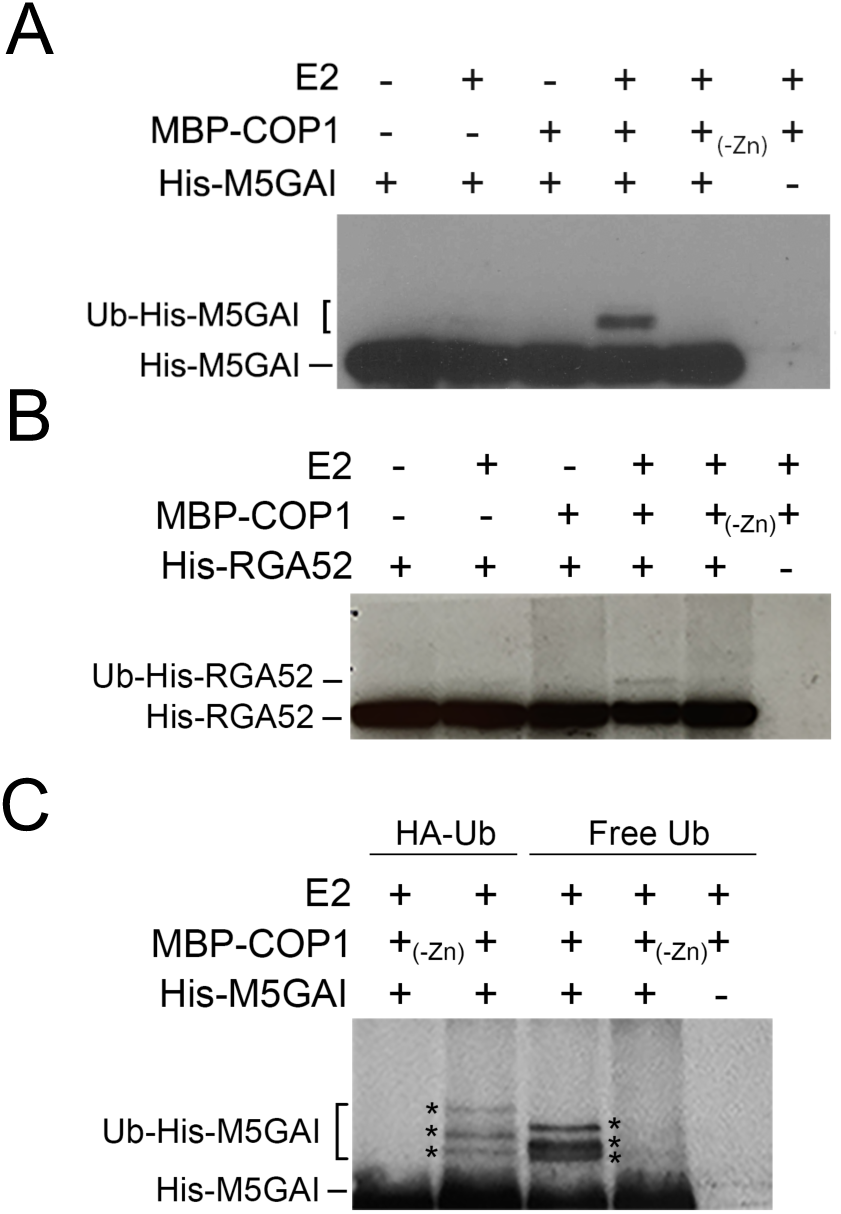
COP1 ubiquitinates GAI and RGA. (A, B) 6xHis-M5GAI (A) and 6xHis-RGA52 (B) ubiquitination assay using recombinant MBP-COP1, rice E2 and unmodified ubiquitin. (C) 6xHis-M5GAI ubiquitination assay using unmodified and HA-tagged ubiquitin. Modified and unmodified 6xHis-M5GAI and 6xHis-RGA52 were detected with anti-GAI and anti-6xHis antibodies, respectively.

### COP1 controls hypocotyl elongation in a DELLA-dependent manner

The growth phenotypes caused by GA deficiency or *cop1* mutations in the dark (30, 34), in response to light cues from neighbors (13, 25) or to warm temperature (10, 14, 35) are very similar. To determine the physiological relevance of the regulation of DELLA levels by COP1, we studied how mutations at *COP1* and *DELLA* genes and GA-treatments impact on the hypocotyl growth rate of dark-grown seedlings transferred to light at 22 °C (de-etiolation) as well as light-grown seedlings transferred to shaded or warm environments. Noteworthy, the patterns differed between the first case, where DELLA levels build up, and the other two cases, where DELLA levels decrease (Fig. 1). In fact, during de-etiolation, growth in the presence of 5 μM GA_4_ promoted the rate of hypocotyl elongation in seedlings transferred to the light but not in seedlings that remained in the dark, suggesting that endogenous GA levels are not limiting in darkness (Fig. 5). As expected, the *cop1* mutants showed reduced growth in darkness; however, they retained a significant growth response to light. This response was only marginally enhanced by adding GAs or by the *gai-td1* and *rga-29* (23) mutations of *DELLA* genes. In other words, during de-etiolation, the rapid inactivation of COP1 does not appear to be rate limiting for the RGA accumulation (Fig. 1) or the growth-inhibition (Fig. 5) responses. Addition of 5 μM GA_4_ promoted growth in light-grown seedlings transferred to shade or to warm temperature (Fig. 5), suggesting that GA signaling is limiting under those conditions. The *cop1-4* and *cop1-6* mutants failed to respond to shade or warm temperatures but the responses were restored both by the application of GAs and by the presence of mutations at *DELLA* genes. This indicates that the responses were limited by the absence of COP1 to target DELLAs to degradation and that reducing the DELLA pool either genetically or via the canonical degradation pathway was enough to rescue the *cop1* phenotype.

**Fig. 5.**
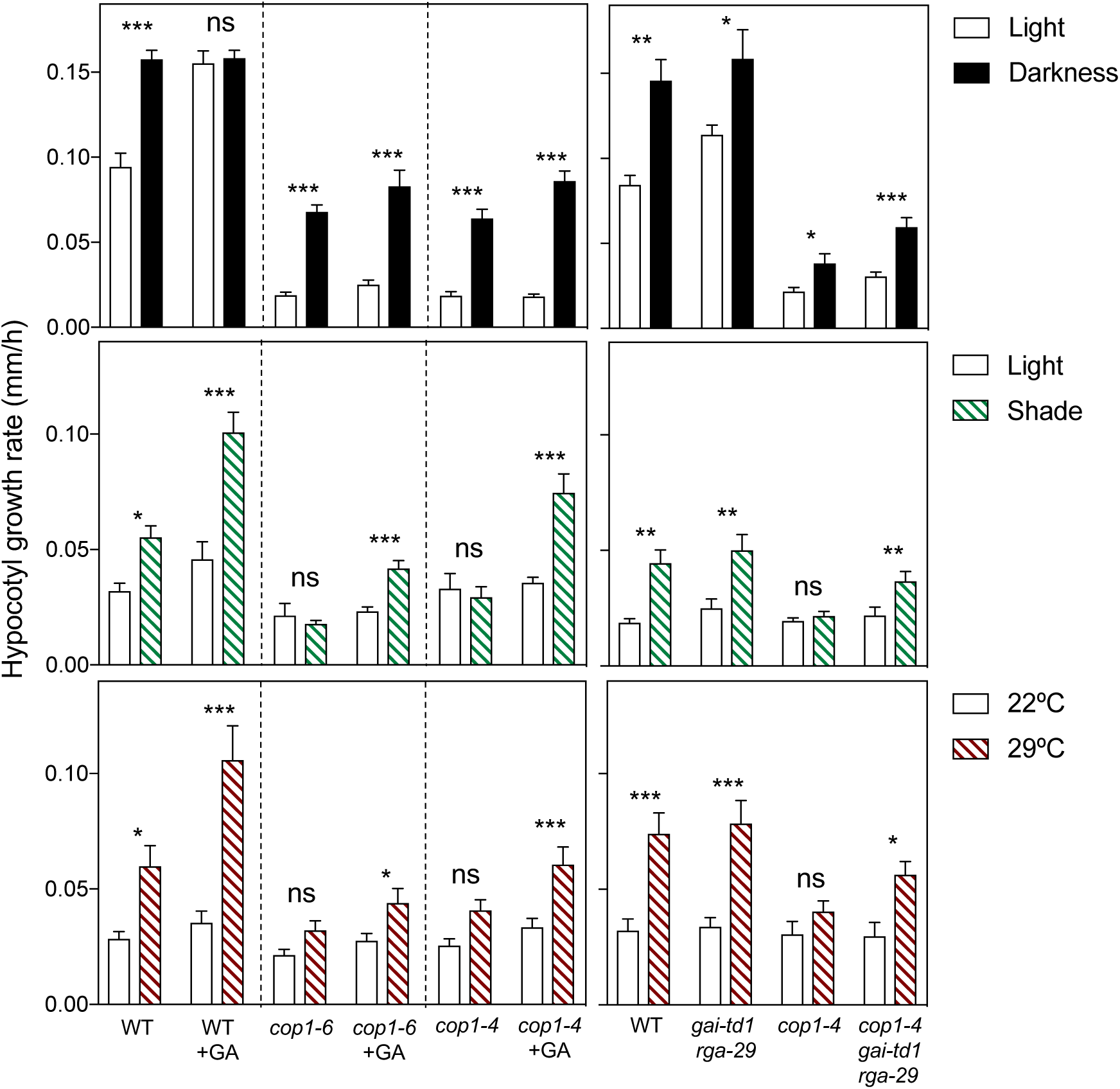
COP1 regulates the rate of hypocotyl elongation in response to shade and warm temperature in a DELLA-dependent manner. Bars indicate the hypocotyl growth rate of seedlings of the indicated genotypes during de-etiolation or after transfer to shade or 29 °C measured over a period of 9 h. Where indicated, seedlings were germinated and grown in the presence of 5 μM GA_4_. Values correspond to the mean and SE of 8 (light treatments) or 24 (temperature treatments) replicate boxes; 10 seedlings were averaged per box replicate. Asterisks indicate that the difference is statistically significant (Student’s *t* test, * *P* < 0.05, ** *P* < 0.01, *** *P* < 0.005; ns, non-significant).

## Discussion

The results presented here establish a functional link between DELLA proteins and COP1, two of the major hubs in the control of plant architecture. The growth of the hypocotyl of *Arabidopsis* shifted from light at moderate temperatures to either warm or shade conditions requires COP1 only if DELLA proteins are present (Fig. 5). These environmental cues reduce phyB activity (2, 36, 37) and enhance COP1 nuclear abundance (Fig. 1*D*), while reducing the levels of RGA in a COP1-dependent manner (Fig. 1*E*). COP1 does not simply reduce DELLA protein abundance by increasing GA levels. First, COP1 migrates to the nucleus and mediates RGA degradation in response to warm temperature well before increasing GA levels (Fig. 1 F-H). Similarly, simulated shade takes more than 4 h to modify GA levels (38) whilst already causing large COP1-mediated effects on RGA at 2 h (Fig. 1*E*). Second, warm temperature or shade reduce the abundance of RGA in the presence of saturating levels of a GA synthesis inhibitor (Fig. 1*A*). Third, warm temperature or shade reduce the abundance of the mutant protein rga-Δ17, which cannot be recognized by GID1 (39) and is fully insensitive to GA (Fig. 1B, C, 2C and *SI Appendix*, Fig. S2). The latter effects require COP1, providing evidence for a branch of COP1 action on DELLA that does not involve activating the canonical GA/GID1 pathway.

COP1 effects on RGA and rga-Δ17 depend on the 26S proteasome (Fig. 2). Taking into account the well-established role of COP1 in E3-ligase complexes that ubiquitinate and target to proteasomal degradation a number of signaling proteins (40), the simplest interpretation of the above observations is that COP1 directly regulates DELLA protein stability. Different results lend support to this hypothesis. First, RGA and GAI interact with COP1 and its complex partner SPA1 in yeast (Fig. 3 A and B) and in planta (Fig. 3C). Second, COP1, SPA1 and GAI or RGA form a tertiary complex in mammalian and plant cells, and this complex is present in nuclear bodies (Fig. 3 D-G and *SI Appendix*, Fig. S3 *A-C* and Fig. S4). Third, COP1 ubiquitinates RGA and GAI in vitro (Fig. 4) and the levels of ubiquitinated RGA *in vivo* are enhanced by COP1 (*SI Appendix*, Fig. S5).

Tight regulation of abundance is a common feature of proteins that act as signaling hubs in mammals (41) and in yeast (42). Post-translational modifications (43–45) and interaction with other transcriptional regulators (46, 47) modulate DELLA activity. However, the mechanism reported here is unique. In contrast to previously reported modes of regulation of DELLA abundance, which converge to control its stability via GA/GID1, the rapid COP1-mediated regulation occurs by a mechanism parallel to the canonical GA/GID1 pathway.

COP1 might represent an ancient regulatory mechanism of control of DELLA levels, preceding the acquisition of the GA/GID1 system because the GA/GID1 system appears in lycophytes (48), whilst orthologs of COP1 and DELLA proteins are already present in the genome of the liverwort *Marchantia polymorpha* (49). Regardless this exciting evolutionary hypothesis, although the GA/GID1 pathway elicits a major control of DELLA levels in vascular plants, co-existence of the COP1 regulation would provide the advantage of a faster adjustment to the sudden fluctuations of the natural environment.

## Materials and Methods

Detailed description of the plant materials and growth conditions, and methods used for protein-protein interaction assays, protein localization and in vitro ubiquitination can be found at *SI Materials and Methods*.

## Acknowledgements

We thank Drs Isabel Lopez-Diaz and Esther Carrera the gibberellin quantification carried out at the Plant Hormone Quantification Service (IBMCP, Valencia, Spain) and Luís López-Molina (University of Geneva, Geneva, Switzerland) for the anti-GAI antibody. We also thank all members of the SIGNAT consortium for helpful discussions about this work. This work was supported by the Spanish Ministry of Economy, Industry and Competitiveness and AEI/FEDER/EU (grants BIO2016-79133-P to DA; BIO2013-46539-R and BIO2016-80551-R to VR), the European Union SIGNAT-Research and Innovation Staff Exchange (grant H2020-MSCA-RISE-2014-644435 to MAB, DA and JJC), the Argentinian Agencia Nacional de Promoción Científica y Tecnológica (grant PICT-2016-1459 to JJC), Universidad de Buenos Aires (grant 20020170100505BA to JJC), the National Institute of General Medical Sciences of the National Institutes of Health (award numbers RO1GM067837 and RO1GM056006 to SAK), by the German Research Foundation (DFG) under Germany’s Excellence Strategy/Initiative (CEPLAS – EXC-2048/1 – Project ID 390686111 to MDZ), by the International Max Planck Research School (IMPRS) of the Max Planck Society, the University of Düsseldorf and Cologne to TB, and NRW-BioSC-FocusLabs CombiCom to NH and MDZ, and MEYS of the Czech Republic (project LQ1601 CEITEC 2020 to BB and MC). NB-T, EI and MG-L were supported by MINECO FPI Program fellowships.

---

**Fig. S1.**
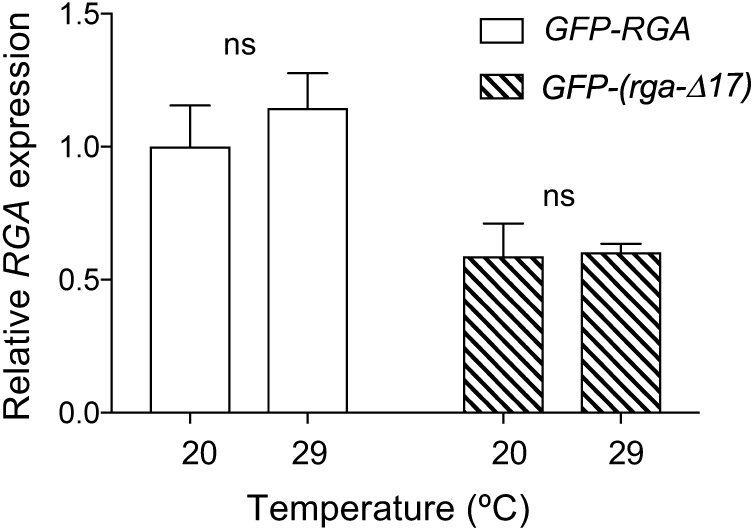
The GA-resistant rga-Δ17 is regulated post-transcriptionally by warm temperature. *RGA* expression in hypocotyls was analyzed by RT-qPCR. Data correspond to mean and standard deviation from three technical replicates. The analysis was performed twice with similar results. ns indicates that the difference is non-significant (Student’s *t* test).

**Fig. S2.**
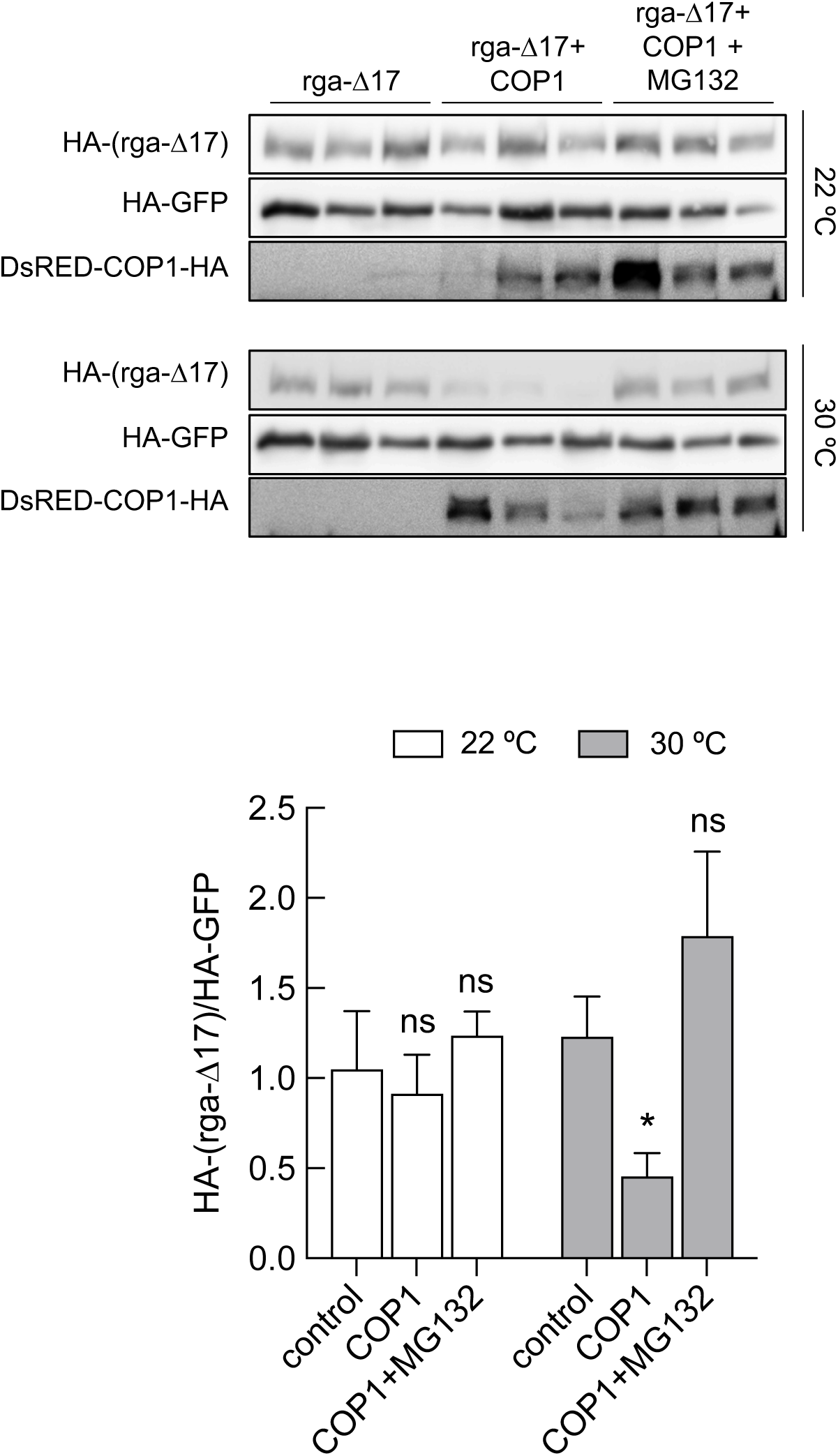
Destabilization of the GA-resistant rga-Δ17 by warm temperature in *N. benthamiana* leaves is dependent on COP1. HA-(rga-Δ17) was transiently expressed alone or with DsRED-COP1-HA in leaves of *N. benthamiana* plants that were transferred to continuous with light at 22 °C or 30 °C for three days. For MG132 treatments leaves were infiltrated with a solution of 25 μM of the inhibitor 8 h before sampling. HA-GFP was used as control to demonstrate the specificity of COP1 action. Blots show data from three individual infiltrated leaves per mixture. Plot shows HA-(rga-Δ17) normalized against HA-GFP. Data are means and SE of 3 leaves from one experiment, repeated twice with similar results. The asterisk indicates that the difference is statistically significant (Student’s *t* test, * *P* < 0.05; ns, non-significant).

**Fig. S3.**
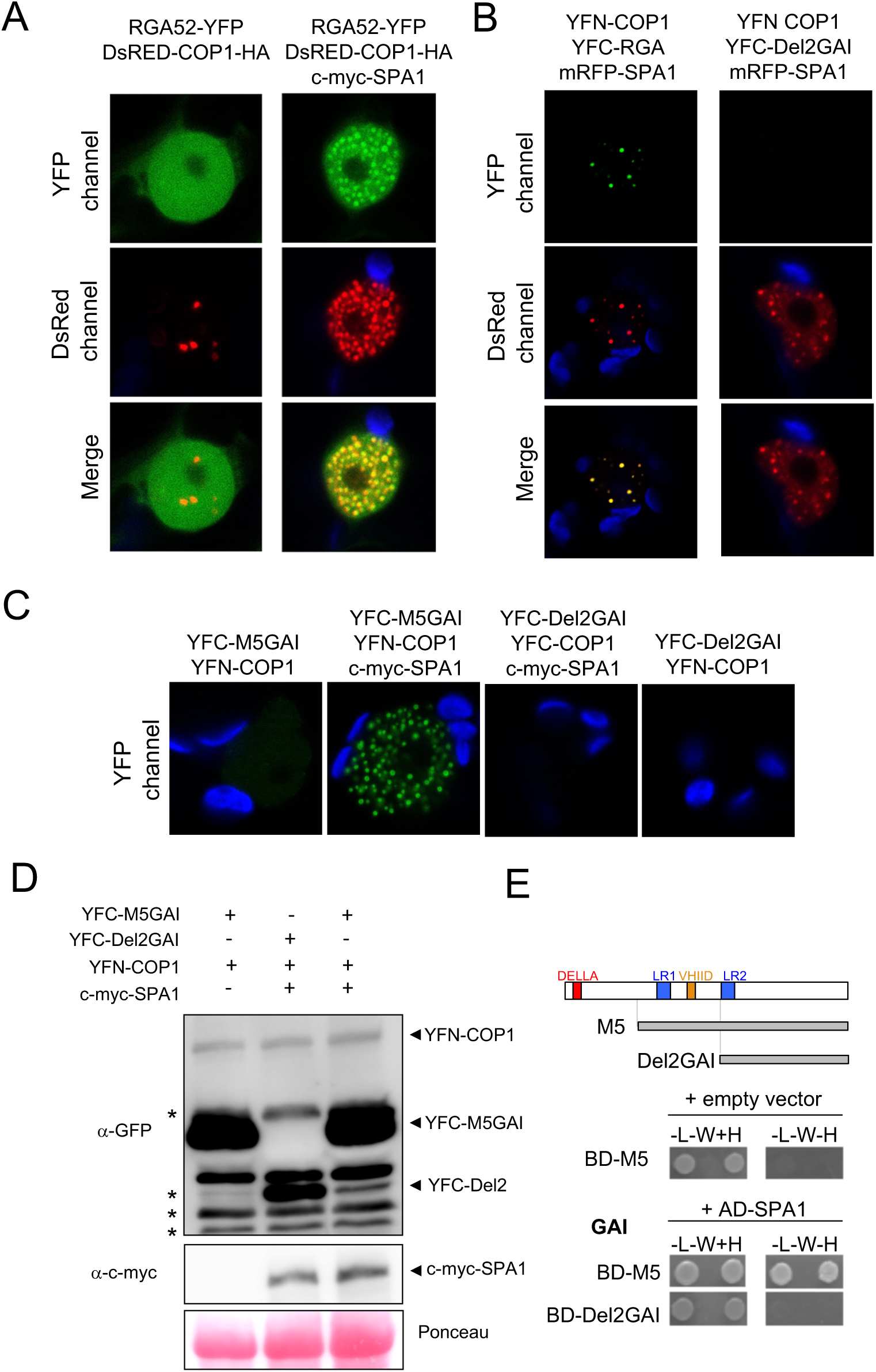
SPA1 helps COP1 recruiting DELLA proteins to nuclear bodies. (A) RGA52-YFP co-localizes with DsRED-COP1-HA in nuclear bodies in the presence of c-myc-SPA1. Fusion proteins were transiently expressed in leaves of *N. benthamiana* and observed by confocal microscopy. One representative nucleus is shown. (B-C) BiFC assay showing that RGA (B) and M5-GAI (C) form a complex with COP1 and SPA1 in nuclear bodies, along with negative controls. The indicated proteins were expressed in leaves of *N. benthamiana* and observed by confocal microscopy. One representative nucleus is shown. (D) Immunoblot analysis of fusion proteins from leaves used for BiFC. The fusion proteins have the expected size: YFN-COP1 (97 KDa), YFC-M5GAI (55 KDa), YFC-Del2GAI (40 KDa) and c-myc-SPA1 (117 KDa). Asterisks indicate non-specific bands, one of them overlapping with YFC-Del2GAI. (E) Y2H assay showing that the GAI deletion Del2GAI does not interact with SPA1.

**Fig. S4.**
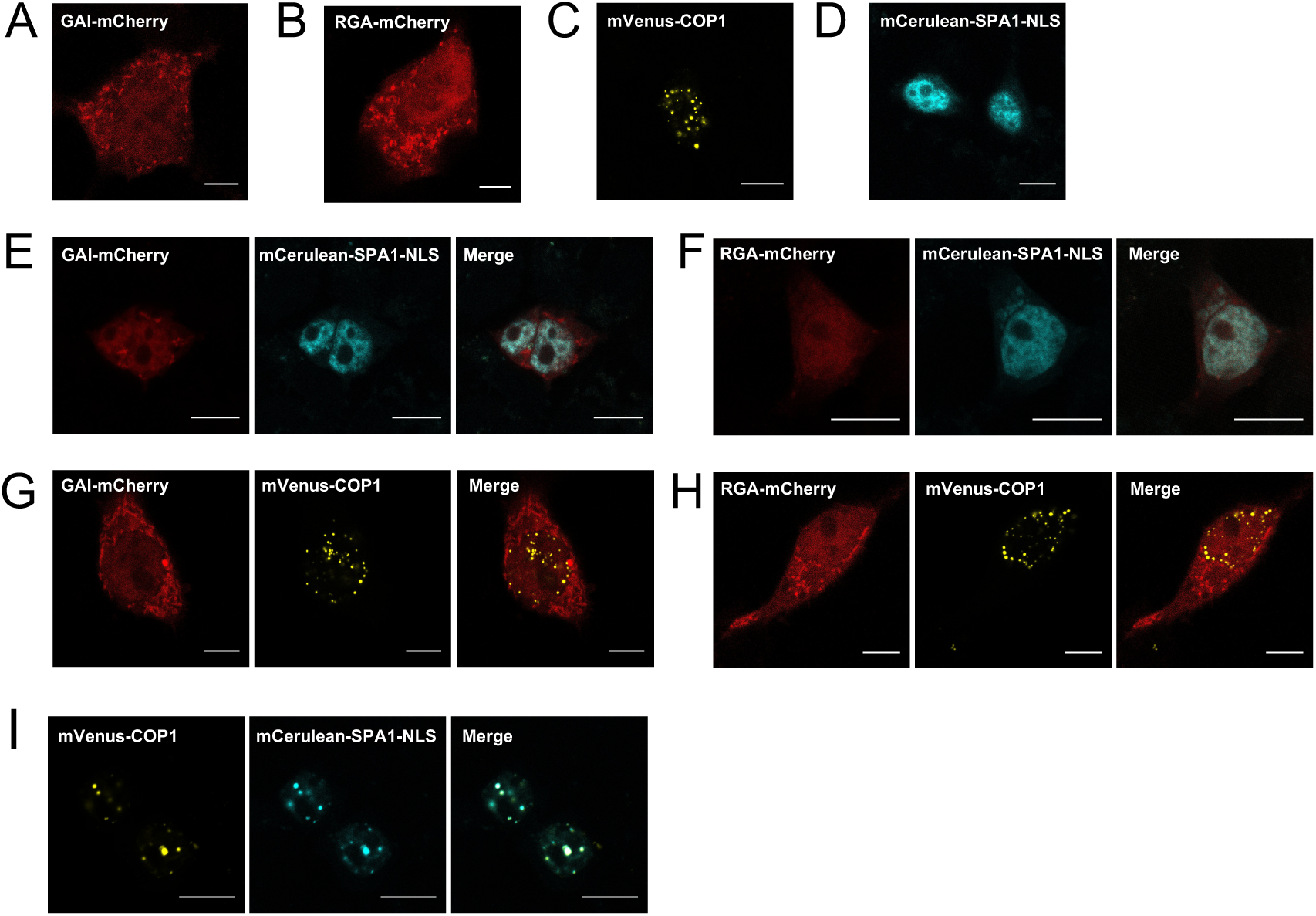
Confocal microscopy analysis of the localization of GAI, RGA, COP1 and SPA1 in animal cells. (A-D) The fusion proteins GAI-mCherry (A), RGA-mCherry (B), mVenus-COP1 (C) and mCerulean-SPA1-NLS (D) were transfected into Human Embryonic Kidney 293T (HEK-293T) cells. GAI and RGA are distributed throughout the whole cell. mVenus-COP1 localizes to nuclear speckle-like structures. mCerulean-SPA1-NLS localizes to the nucleus. (E-I) HEK-293T cells were co-transfected with GAI-mCherry and mCerulean-SPA1-NLS (E), RGA-mCherry and mCerulean-SPA1-NLS (F), GAI-mCherry and mVenus-COP1 (G), RGA-mCherry and mVenus-COP1 (H), or mVenus-COP1 and mCerulean-SPA1-NLS (I). Representative cells are shown. Scale bar represents 10 μm. Note that even when expressed separately, COP1 or SPA1 had weak interactions with DELLAs. For instance, GAI showed more fluorescence inside the COP1 nuclear speckles than outside them (speckles / background fluorescence in G: 1.16 ± 0.02, *P* < 0.05) and inside the nucleus where SPA1 was present than in the cytosol (bright areas in E correspond to the nucleus and in A to the whole cell).

**Fig. S5.**
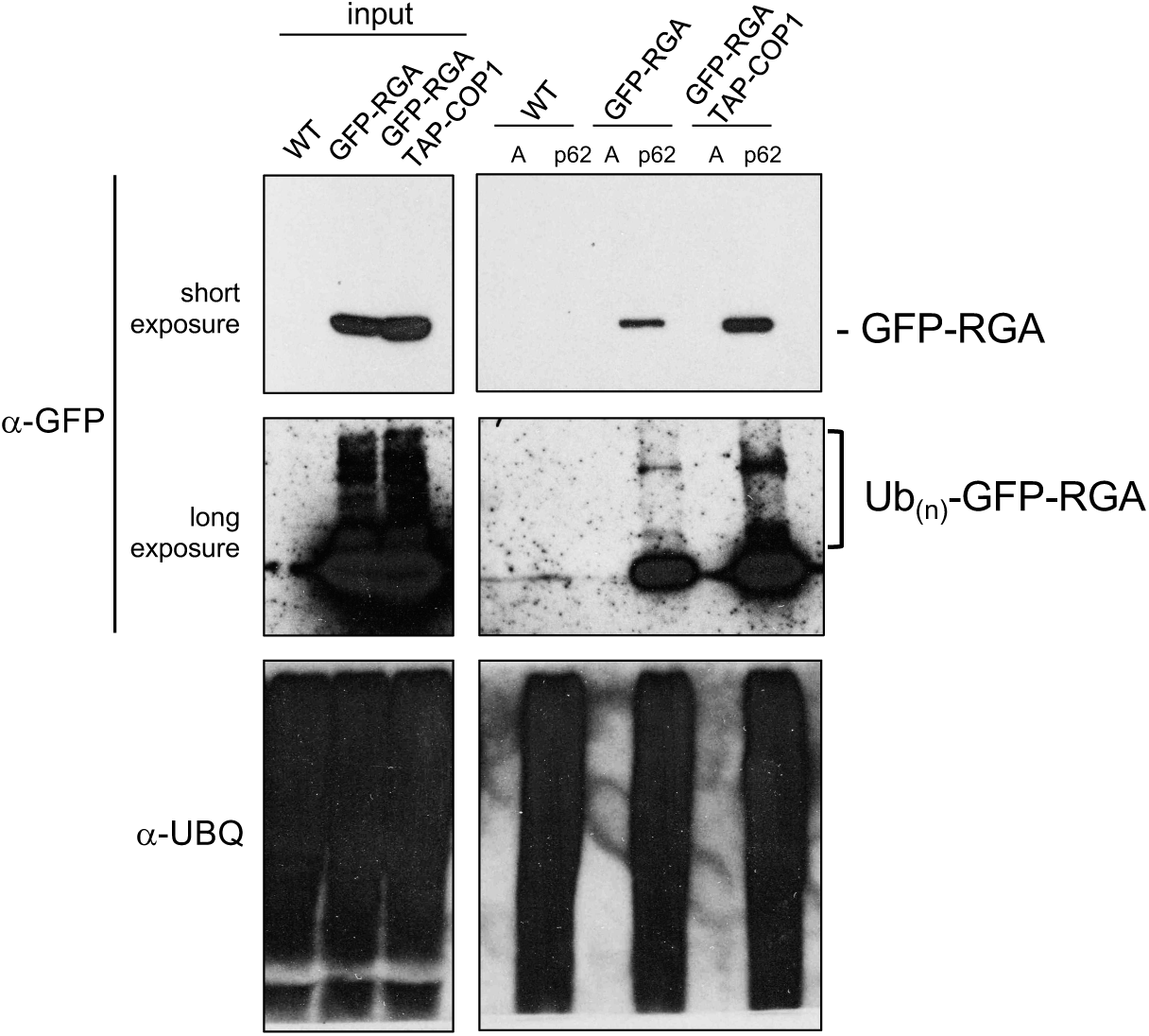
RGA ubiquitination in vivo is enhanced by COP1. *pRGA:GFP-RGA* and *pRGA:GFP-RGA 35S:TAP-COP1* seedlings along with the WT were grown for 5 days at 22 °C and then transferred at 29 °C and treated with 50 μM MG132 for 8 h. Proteins were pulled down with p62 beads or agarose beads (A) as negative control. GFP-RGA and its polyubiquitinated forms were detected with an anti-GFP antibody. Total ubiquitinated proteins were detected with an anti-UBQ antibody.

## Supplementary Methods

### Plant material

Mutants and transgenic lines used in this study that have been described previously: *cop1-4* (1), *cop1-6* (2), *gai-td1* (3), *rga-29* (4), *cop1-4 gai-td1 rga-29* (5), *pRGA:GFP-RGA* (6), *pRGA:GFP-RGA cop1-4* (5)*, 35S:YFP-COP1 cop1-4* (7), and *35S:TAP-COP1* (8). The *35S:TAP-COP1 pRGA:GFP-RGA* double transgenic was prepared by genetic crosses.

### Confocal microscopy in *Arabidopsis*

Seeds were sown in 0.8% agar/water and stratified at 4 °C and darkness for 3-5 days. For the de-etiolation experiments, germination was induced with a 2 h-light treatment. For confocal imaging during de-etiolation, seedlings were grown in darkness for three days and were treated on the fourth day with white light (100 μmol m^-2^ s^-1^, red/far-red=1). For shade treatment, seedlings were grown for 2 days under a 10 h light:14 h dark photoperiod (white light 100 μmol m^-2^ s^-1^, red:far-red= 1). One hour after the beginning of the third day seedlings were shaded with two green filters (LEE #089) following our established protocol in previous shade experiments (9). For the temperature treatment, seedlings were grown for three days at 22 °C in continuous white fluorescent light (25 μmol m^-2^ s^-1^). The fourth day plants were transferred to a chamber at 29 °C with the same lighting conditions, following our established protocol in previous temperature experiments (10). When indicated, either paclobutrazol (a GA biosynthesis inhibitor; final concentration 0.1, 1, 10 or 100 µM) or MG132 (final concentration 100 µM) were respectively added 8 h or 3 h before starting the application of the 29 °C or shade treatments, and confocal fluorescence images of *pRGA:GFP-RGA* were taken 2-4 h after the application of these treatments. In kinetics experiments, confocal fluorescence images of *35S:YFP-COP1 cop1-4* and of *pRGA:GFP-RGA* in the wild-type and *cop1-4* backgrounds were taken at the indicated time points. In experiments including the *pRGA:GFP-rga-Δ17* line, all confocal fluorescence images were taken after 3 day of treatment because saturating levels of the GFP-rga-Δ17 fluorescence signal precluded the detection of fast changes. In some of the experiments including the *pRGA:GFP-rga-Δ17* line, GA_4_ (final concentration 3 µM) was applied to seedlings grown at 22 °C in continuous white fluorescent light (25 µmol m-2 s-1). Confocal fluorescence was obtained with an LSM 5 Pascal Zeiss microscope with a water-immersion objective lens (C–Apochromat 40X/1.2; Zeiss). GFP/YFP were excited with an Argon laser (488nm) and fluorescence was detected at 505-530 nm. Fluorescence intensity was measured using ImageJ. Images were taken from the epidermis and first sub-epidermal cell layers in the upper third part of the hypocotyl. In the figures where shade and 29 °C conditions are compared to a control, data were actually normalized to the corresponding control condition. Chloroplasts autofluorescence was detected between 675 and 760 nm.

### Gibberellin quantification

Seedlings were grown and harvested as described for confocal microscopy experiments with temperature treatments. The ground tissue (about 150 mg of frozen seedlings) was suspended in 80% methanol-1% acetic acid containing internal standards and mixed by shaking for one hour at 4 °C. The extract was kept a −20 °C overnight and then centrifuged and the supernatant dried in a vacuum evaporator. The dry residue was dissolved in 1% acetic acid and passed consecutively through a reverse phase column Oasis HLB (30 mg, Waters) and a cationic exchange Oasis MCX eluted with MeOH, as described elsewhere (11). The final residues were dried and dissolved in 5% acetonitrile-1% acetic acid and the hormones were separated by UHPLC with a reverse Accucore C18 column (2.6 µm, 100 mm length; Thermo Fisher Scientific) with a 2 to 55% acetonitrile gradient containing 0.05% acetic acid, at 400 µL/min over 21 min. The hormones were analyzed with a Q-Exactive mass spectrometer (Orbitrap detector; ThermoFisher Scientific) by targeted Selected Ion Monitoring (tSIM; capillary temperature 300°C, S-lens RF level 70, resolution 70.000) and electrospray ionization (spray voltage 3.0 kV, heater temperature 150°C, sheath gas flow rate 40 µL/min, auxiliary gas flow rate 10 µL/min) in negative mode. The concentration of GA_4_ in the extracts was determined using embedded calibration curves and the Xcalibur 4.0 and TraceFinder 4.1 SP1 programs. The internal standards for quantification were the deuterium-labeled hormones (OlChemim Ltd).

### Real-time-qPCR

*Arabidopsis* seedlings of *pRGA:GFP-RGA* and *pRGA:GFP-(rga-Δ17)* lines were grown for seven days in continuous white fluorescent light (25 μmol m^-2^ s^-1^) at 20 or 29 °C. Hypocotyls were excised and flash frozen in liquid nitrogen. RNA was extracted with the RNAeasy Plant Mini Kit (Qiagen). cDNA was prepared from 1 μg of total RNA with PrimeScript 1^st^ Strand cDNA Synthesis Kit (Takara Bio Inc). PCR was performed in a 7500 Fast Real-Time PCR System (Applied Biosystems) with SYBR premix ExTaq (Tli RNaseH Plus) Rox Plus (Takara Bio Inc). *RGA* was amplified using described primers (12). Expression levels were normalized to *EF1α* (13).

### Degradation assays in *Nicotiana benthamiana*

The *GFP* CDS was cloned into pDONR207 (Thermo-Fisher Scientific) by a BP reaction (Thermo-Fisher Scientific). To prepare an entry vector with *rga-Δ17*, the CDS was amplified by PCR from a cDNA pool obtained from *pRGA:GFP-(rga-Δ17)* seedlings (14) and cloned into pDONR207. The pENTR201-RGA was obtained from the Regia collection (15). The *COP1* CDS was transferred to pEarleyGate202 (16) to create a Flag-COP1 fusion; *GFP*, *RGA*, and *rga-Δ17* CDSs were transferred to pEarleyGate201 to introduce an HA in the N-terminus of each protein; in all cases transfer to the destination vector was done by LR reaction. A pCambia-based binary vector expressing DsRED-COP1-HA under the control of the *35S* promoter was also prepared by GoldenBraid (17). Leaves of one-month-old *N. benthamiana* plants grown under 16 h light:8 h dark photoperiod at 25 °C were infiltrated with the different mixtures of *Agrobacterium tumefaciens* GV3101 cells carrying the vectors and the p19 silencing suppressor. Leaves were harvested at dawn of the fourth day and frozen in liquid nitrogen. For temperature experiments, *N. benthamiana* plants were maintained at 22 °C and continuous white light (LL; 100 μmol m^-2^ s^-1^) or transferred to 30 °C and LL for three days after infiltration. For MG132 treatments leaves were infiltrated with a solution of 25 μM of the inhibitor 8 h before sampling. To determine protein levels, samples were homogenized with 3 volumes of 2x Laemmli buffer and boiled at 95°C for 5 min. Samples were then clarified by centrifugation at room temperature and analyzed by Western blot as described above. Fusion proteins were detected by HRP-conjugated Flag M2 antibody (1:2000; Sigma) and anti-HA-HRP antibody (3F10, 1:2000; Roche). Chemiluminescence was detected with the SuperSignal^TM^ West Pico substrate (Thermo-Fisher Scientific) and imaged with a UVP ChemiDoc imaging system (UVP, LLC). The VisionWorksLS (UVP, LLC) software was used to quantify protein levels.

### Yeast two-hybrid assays

The Y2H, Gateway^TM^-adapted vectors pGBKT7 and pGADT7 were used. The *M5GAI*, *RGA52* and *Del2GAI* CDS cloned in pGBKT7 have been described (18). A pENTR-COP1 vector was prepared by cloning the *COP1* coding sequence (CDS) in the pCR8/GW/TOPO vector (Thermo-Fisher Scientific). Entry vectors for *SPA1*, *SPA2* and *SPA4* were prepared by cloning their CDS in pENTR-3C (Thermo-Fisher Scientific). *COP1*, *SPA1*, *SPA2*, and *SPA4* CDSs were transferred to the pGADT7 vector by an LR reaction (Thermo-Fisher Scientific). The yeast strains Y2HGold and Y187 (Takara Bio Inc) were transformed with the constructs in pGBKT7 and pGADT7 vectors, respectively. Interaction assays were performed in diploid cells obtained by mating and grown in selection media.

### Co-immunoprecipitation

*35S:YFP-M5GAI* and *35S:YFP-RGA52* were prepared by transferring the *M5GAI* and *RGA52* CDSs from pCR8/GW/TOPO-M5GAI and pCR8/GW/TOPO-RGA52 (18) to the pEarleyGate104 vector (16). The CDS of *SPA1* was transferred by LR reaction to the pEarleyGate203 (16) to prepare the fusion c-myc-SPA1. The *35S:DsRED-COP1-HA* construct was used to express recombinant COP1. Leaves of one-month-old *N. benthamiana* plants grown under 16 h light:8 h dark photoperiod at 25 °C were infiltrated with the different mixtures of *A. tumefaciens* C58 cells carrying the vectors and the p19 silencing suppressor. Leaves were infiltrated with a solution containing 50 μM MG132 (Calbiochem) and 10 mM MgCl_2_ 12 h before harvesting. Leaves were harvested at dawn of the fourth day and frozen in liquid nitrogen. Approximately 1 mL of ground, frozen tissue was homogenized in 0.5 mL of extraction buffer (25 mM Tris-HCl pH 7.5, 10% glycerol, 1 mM EDTA pH 8.0, 150 mM NaCl, and 1x protease inhibitor cocktail [cOmplete, EDTA-free; Roche]). Extracts were kept on ice for 15 min and cell debris was removed by centrifugation at maximum speed in a benchtop centrifuge at 4 °C twice. Total proteins were quantified by Bradford assay. One hundred and sixty μg of total proteins were denatured in Laemmli buffer and set aside to be used as input. Eight hundred μg of total protein in 1 mL of extraction buffer were incubated with 50 μL of anti-GFP-coated paramagnetic beads (Miltenyi) at 4 °C for 2 h in a rotating wheel. Extracts were loaded into μColumns (Miltenyi) at room temperature. Columns were washed four times with 200 μL of cold extraction buffer and proteins were eluted under denaturing conditions in 70 μL of elution buffer following manufacturer instructions. Sixty-three μL of the immunoprecipitated samples (90%) were loaded in a 12% SDS-PAGE along with 60 μg of input. Proteins were transferred to a PVDF membrane and sequentially immunodetected with an anti-HA-HRP antibody (3F10, 1:5000; Roche) and, after strip-out, with an anti-c-myc antibody (9E10, 1:1000; Roche). The remaining 10% of the immunoprecipitated samples along with 60 μg of input were processed in the same way but immunodetected with an anti-GFP antibody (JL8, 1:5000; Clontech). Chemiluminiscence was detected with SuperSignal^TM^ West Femto (Thermo-Fisher Scientific) and imaged with the LAS-3000 imager (Fujifilm).

### Co-localization assays in *N. benthamiana*

pCambia-based binary vectors expressing RGA52-YFP alone or with DsRED-COP1-HA under the control of the *35S* promoter were prepared by GoldenBraid (17). *35S:YFP-GAI* was described previously (19). The *RGA* CDS was transferred by LR reaction to pEarleyGate104 to prepare a YFP-RGA fusion. Leaves of one-month-old *N. benthamiana* plants grown under 16 h light:8 h dark photoperiod at 25 °C were infiltrated with the different mixtures of *A. tumefaciens* C58 cells carrying the vectors and the p19 silencing suppressor. Leaves were infiltrated with a solution containing 50 μM MG132 and 10 mM MgCl_2_ 12-16 h before imaging by confocal microscopy (Zeiss 780 Axio Observer) with a water-immersion objective lens (C-Apochromat 40X/1.2; Zeiss) the fourth day after infiltration. YFP was excited with an Argon laser (488 nm) and detected at 520-560 nm. DsRED was excited the DPSS 561-10 laser (561 nm) and detected at 580-650 nm. Samples were kept in darkness on the day of imaging. Fluorescence from YFP was detected between 520-560 nm, and fluorescence from DsRED between 580 and 650 nm. Chloroplasts autofluorescence was detected between 675 and 760 nm.

### Bi-molecular fluorescence complementation

The pCR8/GW/TOPO-del2GAI has been described (18). The *M5GAI, RGA52, RGA*, and *Del2GAI* CDSs were transferred to the pMDC43-YFC (20) vector, while the *COP1* CDS was transferred to pMDC43-YFN (20). The CDS of *SPA1* was transferred by LR reaction to the pUBQ10-mRFP (21) to prepare the fusion mRFP-SPA1. Leaves were infiltrated, treated and imaged as described for co-localization assays. *A. tumefaciens* cells carrying the p19 silencing suppressor were not included in the mixtures. YFP fluorescence was detected as described above. mRFP fluorescence was detected as described for DsRED.

### Co-localization assays in mammalian cells

The list of oligonucleotides and details for vector construction are in Supplemental Tables 1 and 2, respectively. Human embryonic kidney cells (HEK-293T, ATCC CRL-11268), were maintained in Dulbecco’s modified Eagle’s medium (PAN, cat. no. P04-03550) supplemented with 10% fetal calf serum (FCS, PAN, cat. no. P30-3602) and 1% penicillin/streptomycin (PAN, cat. no. P06-07100). For confocal imaging, cells were seeded onto glass coverslips placed in cell culture wells. For transfection, 40,000 cells per well of a 24-well plate, were transfected using polyethylenimine (PEI, linear, MW: 25 kDa, Polyscience) as described elsewhere (22). The medium was exchanged 5 h post transfection. In co-transfections, all plasmids were transfected in equal amounts (weight-based). For confocal imaging, cells on glass coverslips were fixed with 4% PFA for 10 min on ice followed by 10 min at room temperature. Subsequently, cells were washed once with ice-cold PBS. Coverslips were embedded in Mowiol 4−88 (Roth) containing 15 mg mL^-1^ 1,4-diazabicyclo[2.2.2]octane (DABCO, Roth) and mounted onto glass microscope slides as described (23). Cells were imaged with a confocal microscope (Nikon Instruments Eclipse Ti with a C2plus confocal laser scanner, 60× oil objective, NA = 1.40). mCherry, mVenus and mCerulean were visualized using excitation lasers of 561, 488, 405 nm and emission filters of 570−620, 535-550, 425−475 nm, respectively.

### Analysis of confocal images of mammalian cells

Image acquisition, analysis and processing were performed with Fiji. The workflow for data analysis is as follows. The mVenus-COP1 channel image was duplicated and adjusted with a low threshold to only mark speckles as regions of interest. Then, the measurement was set to redirect to the DELLA-mCherry channel image. Particles of a size between 100-15.000 square pixels and of any shape were analyzed for their mean fluorescence intensity given in absolute gray values. Additionally, 5 random areas outside of the speckles in the nucleus of each cell were analyzed for their mean fluorescence intensity in the DELLA channel and the mean background fluorescence intensity was calculated. The ratio (DELLA speckle – DELLA background)/DELLA background was calculated for each speckle with the respective background. The mean ratio per cell was calculated. Outlier determination was performed using the quartile method, i.e. values smaller than Q1-1.5(Q3-Q1) or larger than Q3+1.5(Q3-Q1) were excluded in each condition (with or without mCerulean-SPA1-NLS).

### Pull-down assays with p62-beads

Wild-type Col-0, *pRGA:GFP-RGA* and *pRGA:GFP-RGA 35S:TAP-COP1* seedlings were grown for five days at 22 °C in continuous white light (25 μmol m^-2^ s^-1^) and then temperature was shifted to 29°C for 8 h under the same lighting condition. Seedlings were treated with 50 μM MG132 during the high-temperature treatment. Proteins were extracted in buffer BI [50 mM Tris-HCl, pH 7.5, 20 mM NaCl, 0.1% Nonidet P-40, and 5 mM ATP, 1 mM PMSF, 50 μM MG132, 10 nM Ub-aldehyde, 10 mM N-ethylmaleimide, and plant protease inhibitor cocktail (Sigma-Aldrich)] before incubation with pre-washed p62 agarose (Enzo Life Sciences) or the agarose alone at 4 °C for 4 h. Beads were washed twice in BI buffer and once with BI buffer supplemented with 200 mM NaCl. Proteins were eluted in Laemmli buffer at 100 °C. The eluted proteins were separated by SDS-PAGE and analyzed by immunoblotting using anti-Ub (1:1000; Enzo Life Sciences) or anti-GFP antibodies (JL-8, 1:5000; Clontech).

### In vitro ubiquitination assay

Assays were performed as previously reported (24) with minor modifications. Ubiquitination reaction mixtures contained 50 ng yeast E1 (Boston Biochem), 50 ng rice 6xHis-Rad6 E2 (25), 10 μg HA-labelled or 10 μg unlabelled ubiquitin (Boston Biochem), and 2 μg MBP-COP1 (previously incubated with 20 μM ZnCl_2_) in 30 μL of reaction buffer (50 mM Tris pH 7.5, 5 mM MgCl_2_, 2 mM ATP and 0.5 mM DTT). As a substrate, 50 ng of 6xHis-M5GAI or 6xHis-RGA52 fusions were used per reaction. After 2 h incubation at 30 °C, reaction mixtures were stopped by adding 30 μL of Laemmli buffer, and a half of the mixtures (30 μL) were boiled for 5 min and separated by 7.5% SDS-PAGE. 6xHis-M5GAI and 6xHis-RGA52 were detected using anti-GAI (26) (1:10000) and anti-6xHis (Sigma) antibodies, respectively.

### Growth assays

Seeds were sown in 0.8% agar/water with or without 5 μM GA_4_ and stratified at 4 °C and darkness for 3-5 days. For the de-etiolation experiments, germination was induced with a 2 h-light treatment. On the day of treatment plants were imaged with a digital camera before the beginning and after the end of the 9 h-treatment and hypocotyl elongation during this period was measured with image processing software (9). Treatments were as described for confocal imaging. Seedlings kept in darkness, unshaded or at 22 °C were used as controls of the respective treatments.

---

**Supplemental Table 1.**
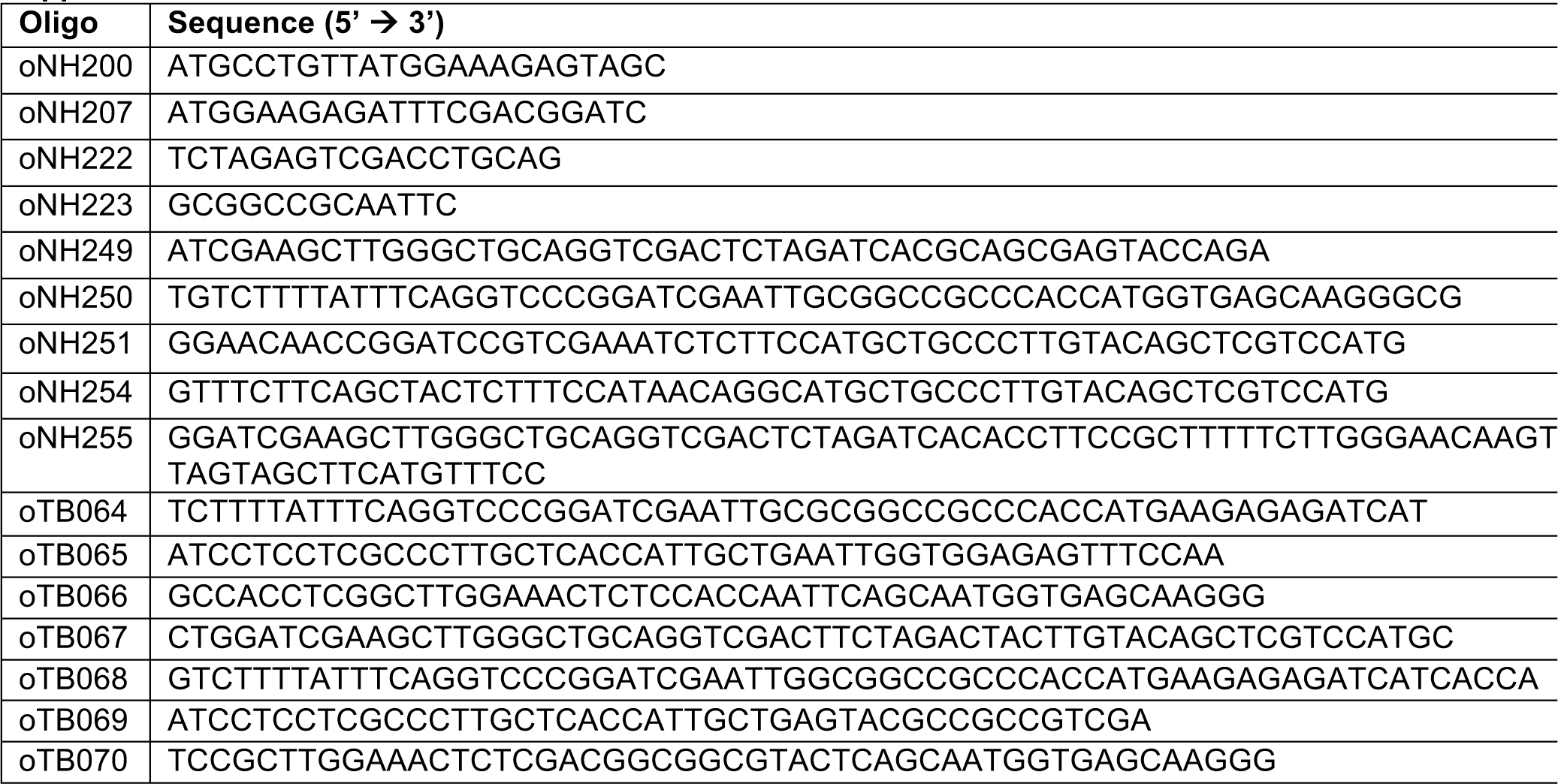

**Supplemental Table 2.**
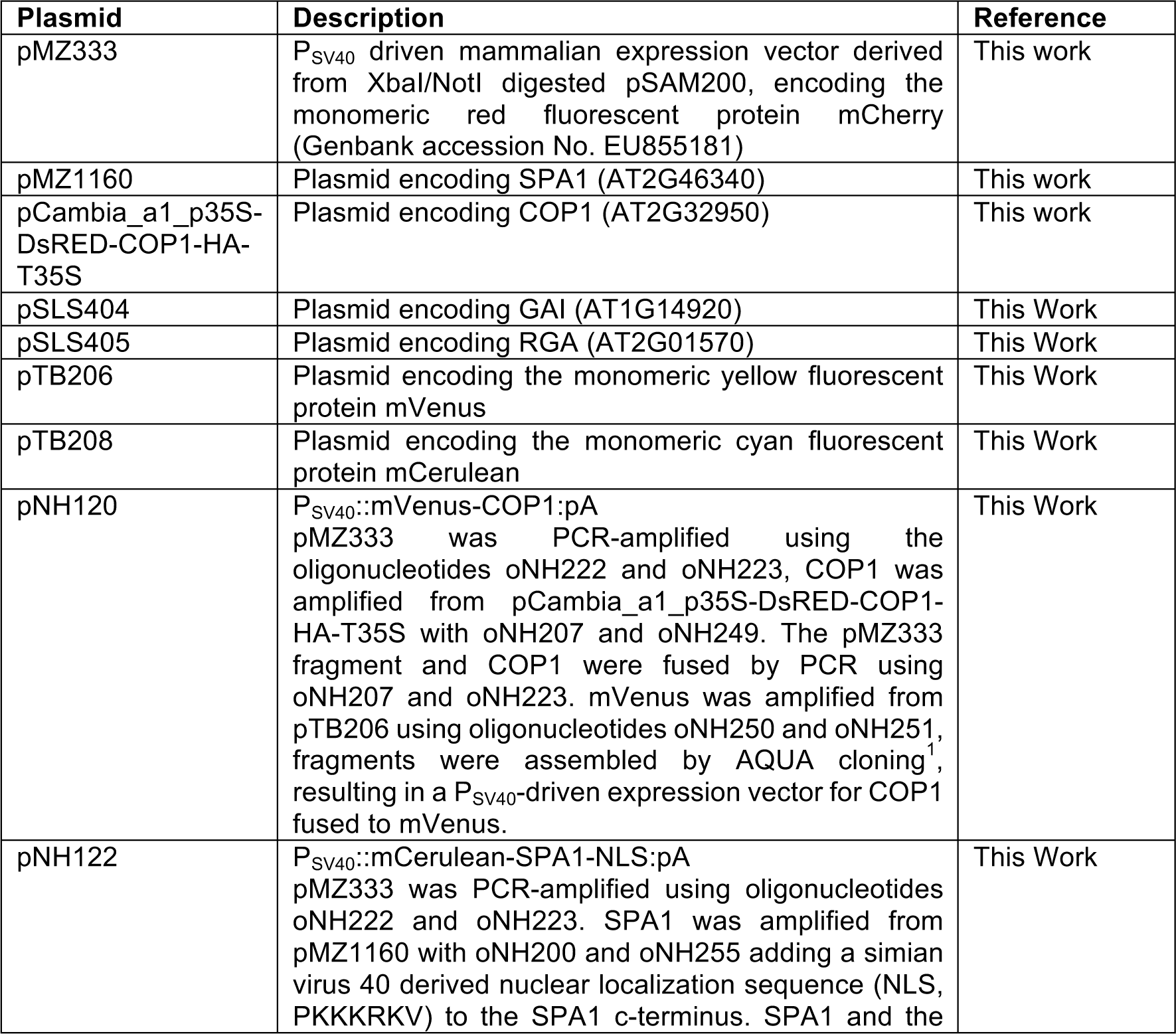
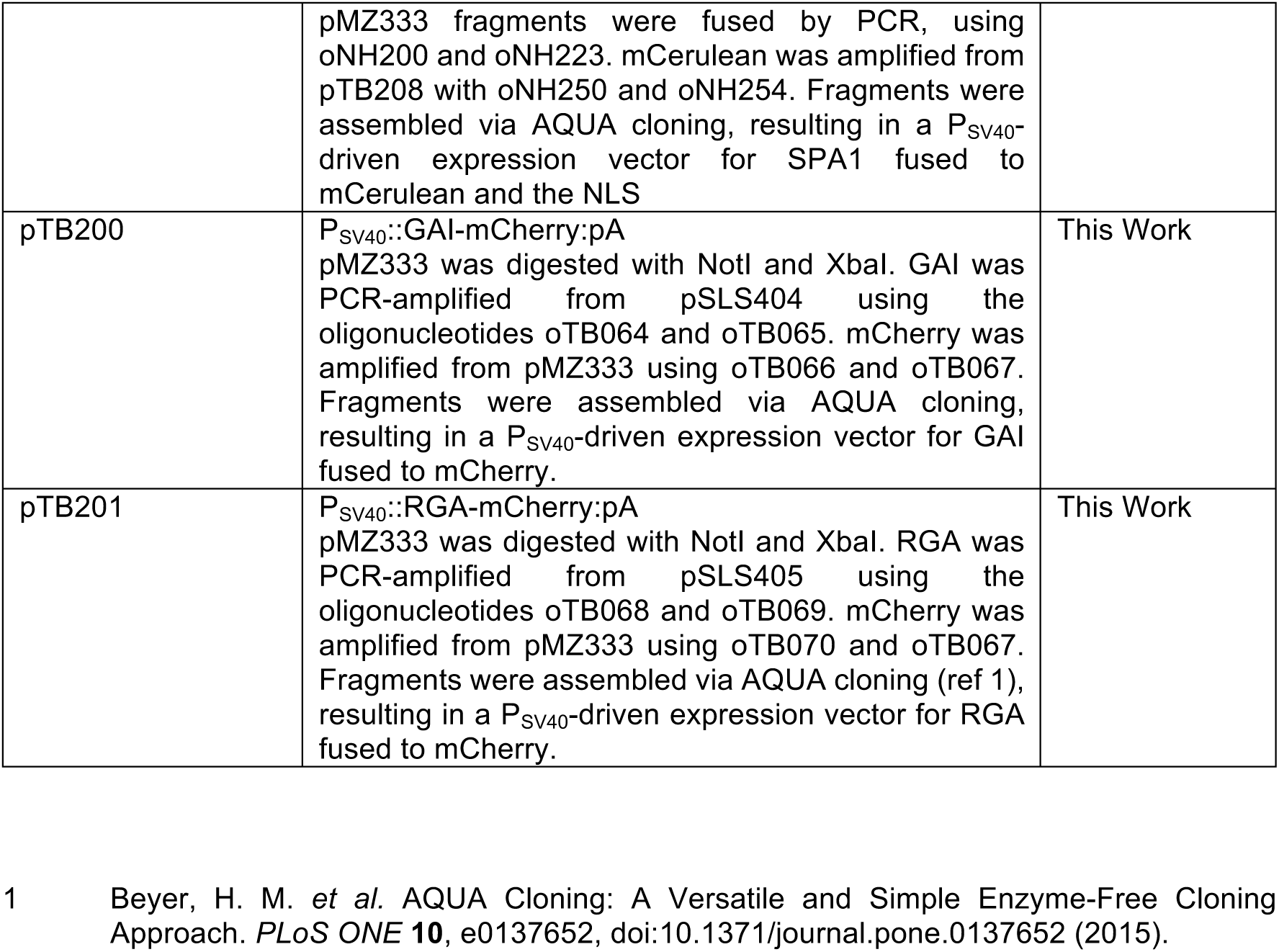

